# The long-standing relationship between replication timing, gene expression, and chromatin accessibility is maintained in early mouse embryogenesis

**DOI:** 10.1101/2025.11.10.687227

**Authors:** Juan Carlos Rivera-Mulia

## Abstract

Replication timing (RT) is the temporal program of genome duplication that is coordinated with 3D genome organization and gene expression^1,2^. How and when RT is established during mammalian development are questions that remain unsolved. Four recent studies provided the first insights; however, they reached to contradictory conclusions. Nakatani et al. reported a poorly defined RT in mouse zygotes that is progressively consolidated by the 4-cell stage^3^; Takahashi et al. and Halliwell, et al. reported a complete RT absence in the zygote with abrupt emergence at later stages^4,5^; finally, Xu et al.^6^ reported a RT already established in the zygote. Discrepancies might reflect differences in computational methods and thresholds applied to sparse single-cell RT data. More recently, Shetty et al. concluded that RT is established in the zygote, and that transcriptional activation and chromatin accessibility emerge at late replicating regions^7^. Their conclusions challenge the long-standing relationship between gene expression and early replication^1,2,8–10^ and contradict previous studies^3–6^. This critique demonstrates that methodological flaws and artifacts, stemming from inadequate sample acquisition and data processing led to unreliable RT inference, rendering the conclusions on RT establishment and its inverse correlation with gene expression erroneous. Moreover, re-analysis of their data, with stage-matched controls from previous studies^3,5^, demonstrates that the canonical relationship between RT, transcriptional activity, and chromatin accessibility are indeed still standing during the early mammalian embryogenesis.

## Results and discussion

### S-phase representation and RT mapping reliability

A recent publication by Shetty et al. analyzed the RT establishment during early mouse embryogenesis and its links to the emergence of the transcriptome and chromatin accessibility. They concluded that *“RT is established at the 1-cell stage prior to zygotic gene activation (ZGA)”*. However, their data do not reflect the early embryogenesis RT. scRepli-seq analyses require entire S-phase representation^11–13^. Thus, previous studies relied on: 1) exhaustive sampling at hourly intervals; 2) large embryo numbers (≥500); 3) replication validation by EdU staining; and 4) multiple replicates, including *in vivo* and *in vitro-* generated embryos, parthenogenetic zygotes, and isolated maternal and paternal pronuclei^3–6^. In stark contrast, Shetty et al. is restricted to ∼40 cells per stage ((Zygote n=36, 2-cell n=42, 4-cell n=43), a single replicate, a unique time point, and without any validation that the collected cells were indeed in S-phase. These limitations hindered precise RT profiling. Although authors state that can map RT and transcriptomes from as low as 10 cells, benchmarking and down-sampling analyses are missing. By reanalyzing their data including replicated genome percentages (standard in scRepliseq^3–5,11,12^), I demonstrate significant S-phase representation gaps. There is an absence of early/middle S-phase cells in the zygote and 4-cell stages, and middle/late S-phase in the 2-cell stage (**Fig. 1a**). This incomplete S-phase representation introduces artifacts and leads to unreliable RT profilong^11–13^; thus, explaining their low resolution and flat RT profiles (**Fig. 1b**). In fact, their own diagnostic analyses based on *Twidth* values (the time for a genomic region to be replicated in the population) yielded unusually high values (>3.0), which is inconsistent with the expected range of 1.5-2.7 in robust, reliable scRepli-seq datasets^11,13^, thus indicating unsuccessful RT profiling. Authors reported that *“70% of total bins across the genome were conserved”* relative to previous studies; however, this was restricted to zygotes and is lacking formal cross-study comparisons (stage-specific concordance, genome-wide correlation analyses, or statistical significance of agreement). Thus, to evaluate the similarity degree, I compared their RT profiles to previous reports focusing on the 4-cell stage, where all previous studies agree^3–6^. This comparison corroborated high concordance between previous studies. However, data from Shetty et al.^7^ differ substantially (**Fig. 1c**), which is inconsistent with the reported 72% overlap with Nakatani et al^3^, undermining the reliability of the RT metrics and conclusions derived by Shetty et al. Together, these observations indicate that the data in this study do not reliably capture RT.

**Figure 1.**
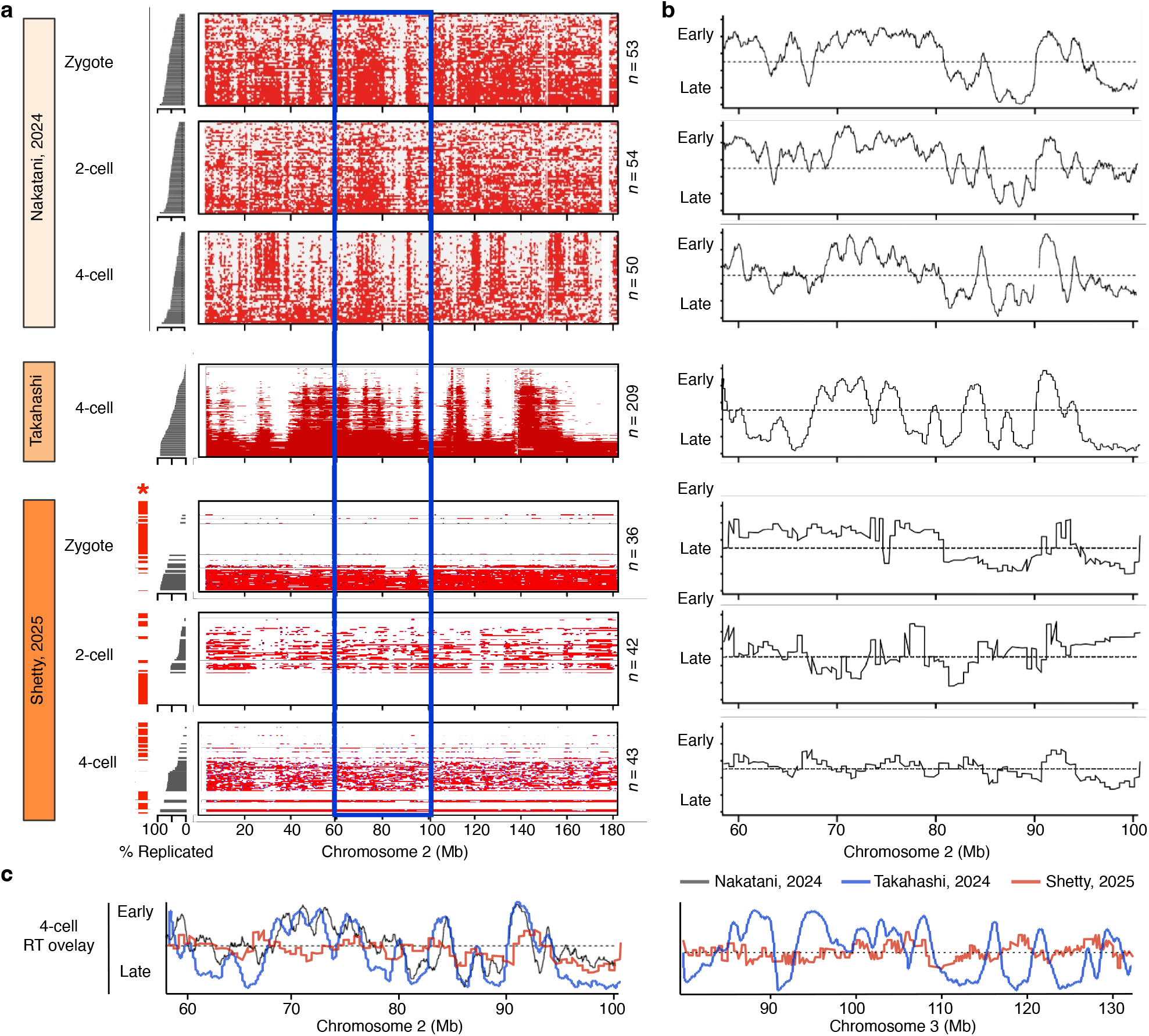
S-phase representation and RT programs from early mouse embryogenesis across studies. **a**, Single cell RT data from zygotes, 2-cell, and 4-cell stages. Datasets were obtained from Nakatani et al.^3^; Takahashi et al.^5^; and Shetty et al.^7^. Cells were ranked based on their percentage of replicated genome (side bar plots). Significant S-phase representation gaps are shown as red bars. Binarized single-cell RT heatmaps are shown with each row representing a single-cell binarized RT (red= replicated, white=unreplicated). Sample sizes are indicated next to the heatmaps. **b**, pseudo-bulk RT profiles for the region highlighted with the blue rectangle in (**a**). **c**, overlay of pseudo-bulk RT profiles at the 4-cell embryo stage derived from the distinct studies.

Multiple assertions by Shetty et al. on RT regulation, such as the decrease in replication domain sizes, and RT changes during early embryogenesis are also made without corresponding quantitative analyses. In fact, core RT features, such as domain sizes, initiation zones, or timing transition regions were not identified, and genomic fractions of preserved versus RT shifting quantification, or RT domain consolidation analyses are missing. Overall, the presented data is insufficient to support their conclusions.

### Transcriptome and Gene Expression analyses

Transcriptome analyses conducted by Shetty et al. also requires scrutiny. The accuracy of stage-specific expression is potentially confounded by maternally inherited RNAs, and their analyses lack critical cross-study validation for post-ZGA genes across all stages, batch correction, and gene-by-gene correspondence analyses. Moreover, reported ontology enrichments are not relevant to preimplantation biology. Furthermore, reanalysis of their scRNA-seq data shows that several genes identified as *“top differentially expressed genes”* and *“stage-specific”* are neither significantly changing nor truly stage-specific (**Fig. 2a**). For instance, *Rif1* and *Orc1-6* are essential for early embryogenesis and then, expected to be expressed. The maternally inherited factor *Npm2*, reported to be drastically downregulated in 2-cell and 4-cell stages, did not show significant expression changes upon re-analysis. Likewise, the major ZGA regulator, *Dux*, identified by Shetty et al. as 2-cell stage-specific, was not detected in their data upon re-analysis (**Fig. 2a**). These inconsistencies indicate that the assertion that *“transcriptome profiles recapitulated stage-specific expression”* requires further validation.

**Figure 2.**
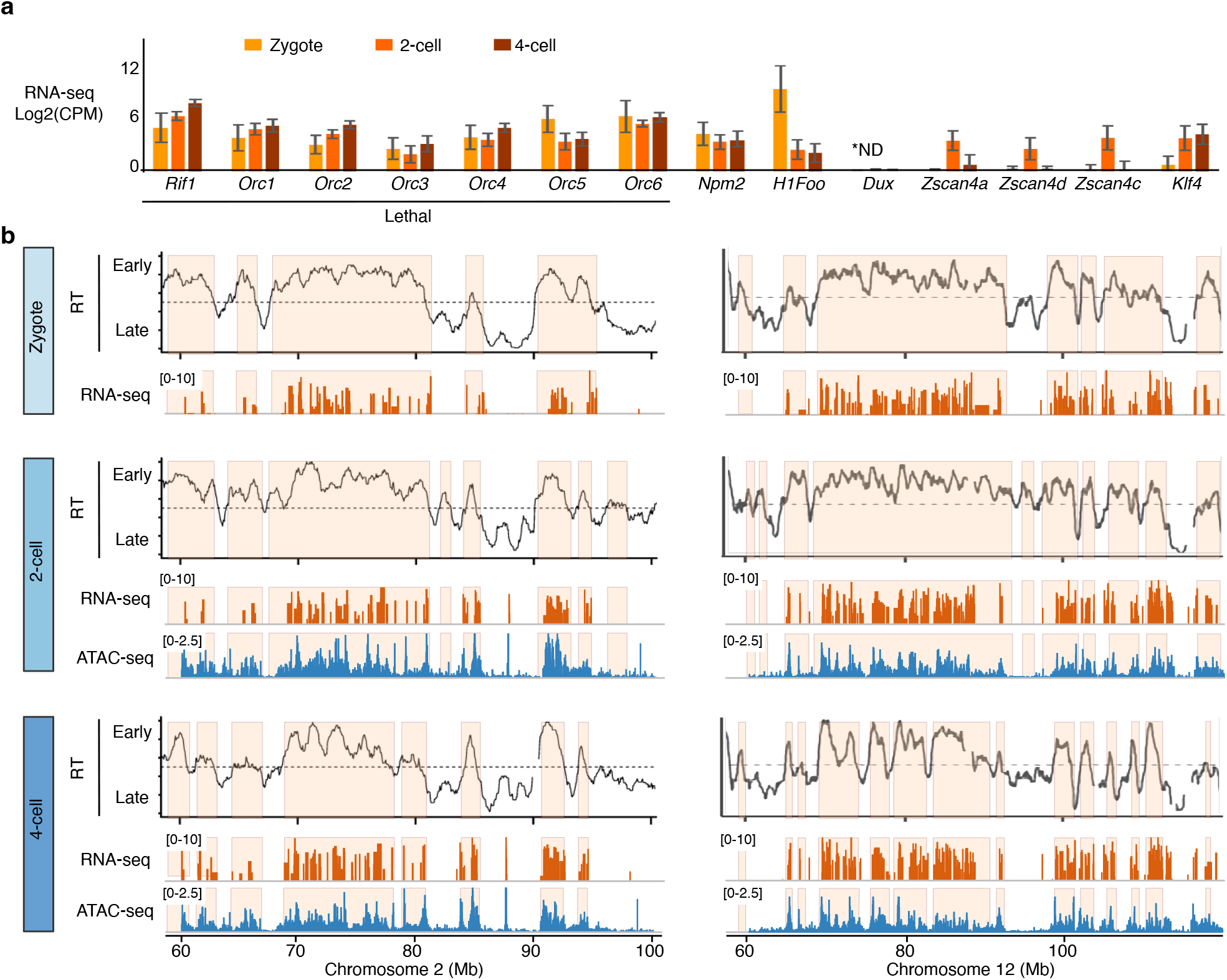
RT correlates with transcriptional activity and chromatin accessibility in early mouse embryogenesis. **a**, Gene expression patterns from selected genes in Shetty et al.^7^. Median Log2(CPM) values and standard deviation from scRNA-seq are shown. *ND=not detected. **b**, RT profiles (pseudo-bulk RT), RNA-seq, and ATAC-seq genomic tracks are shown from zygotes, 2-cell, and 4-cell stages. RT data was obtained from Nakatani et al.^3^; RNA-seq data from Shetty et al.^7^; ATAC-seq data from Wu et al.^14^. Early replicating regions are highlighted in each genomic track.

### Chromatin accessibility analyses

Integrative analyses of RT and chromatin accessibility by Shetty et al. are also questionable, not only because the unreliable RT measurements but also given the unclear sources of reference ATAC-seq data. Authors state that the negative correlation between RT and open chromatin was derived from comparisons of their RT data against reference ATAC-seq data^14^. However, the reference data used for this analysis does not include data from zygotes due to technical limitations as discussed in the original publication^14^. It is therefore unclear how these values were obtained in Shetty et al., and how they derived their conclusions on the relationship between RT and chromatin accessibility.

### RT links to transcriptional activity and chromatin accessibility

The most striking and problematic conclusion drawn by Shetty et al. is that late replicating regions correlate with higher gene expression and open chromatin in early embryos. Authors claim that RT is already established in the zygote, and that transcriptional induction emerges at later replicating regions that have more open chromatin. These claims directly conflict with the foundational knowledge of genome organization and RT in somatic and pluripotent cells, where early replication is strongly associated with actively transcribed, accessible chromatin. Hence, to evaluate these relationships with accuracy, I compared the RT profiles from zygotes through the 4-cell embryonic stage using reliable publicly available RT data^3^, with the corresponding scRNA-seq and ATAC-seq datasets analyzed in Shetty et al.^7^. In contrast with the conclusions from Shetty et al., and consistent with the observations from multiple somatic cell types and across species, these results demonstrate that gene expression is closely associated with early replicating regions from the zygote through the 4-cell stage (**Fig. 2b**). Similarly, ATAC-seq peaks, indicative of open chromatin/accessibility, are closely associated with early replicating regions when analyzed using robust reliable RT data (**Fig. 2b**).

## Conclusions

The core conclusions by Shetty et al. —that RT is established prior to ZGA, and that transcriptional induction and open chromatin emerge preferentially at late replicating regions—are not supported by their data. This critique demonstrates that their conclusions can be attributed to artifactual RT data stemming from methodological limitations. Moreover, the re-analysis of their data, paired with reliable RT data (Nakatani et al.^3^) and ATAC-seq data (Wu et al.^14^) consistently demonstrates that both gene expression and chromatin accessibility emerge at regions associated with early replication. Therefore, the canonical relationship between RT, transcriptional activity, and chromatin accessibility is maintained during early mammalian embryogenesis.

## Methods

Re-analyses presented in this critique rely on publicly deposited sequencing data and published results. Public data were sourced from NCBI Gene Expression Omnibus (GEO) with accession numbers:

- scRepli-seq data from Shetty et al.^7^, GEO accession GSE284010.
- scRNA-seq data from Shetty et al.^7^, GEO accession GSE284010.
- scRepli-seq data from Nakatani et al.^3^, GEO accession GSE218365.
- scRepli-seq data from Takahashi et al.^5^, GEO accession GSE255458.
- ATAC-seq data from Wu et al.^14^, GEO accession GSE66581.
- Other contextual data were drawn from studies including Halliwell et al.^4^ and Xu et al.^6^.

### Binarized scRT

Binarized values for each developmental stage were obtained from the GEO repository (GSE284010, GSE218365, GSE255458). Binarized RT values per cell were ranked based on the percentage of replicated DNA. S-phase representation was derived from the original publications from Nakatani et al.^3^ and Takahashi et al.^5^. For Shetty et al., S-phase progression was divided into 60 bins and cells ranked by their percentage of replicated DNA and empty bins were identified (S-phase gaps).

### Average RT profiles

Aggregated RT profiles were derived from the original publications from Nakatani et al.^3^ and Takahashi et al.^5^, and Shetty et al.^7^ (GSE284010, GSE218365, GSE255458).

### Gene Expression

Transcriptome data was obtained from GSE284010. Median Log2(CPM) values and standard deviations were calculated per developmental stage.

### Chromatin accessibility

ATAC-seq data was obtained from GSE66581. MACS-called peaks were visualized per developmental stage.

## Author Contributions

J.C.R.-M. conceived the critique, performed the re-analysis of publicly available datasets, generated all figures, and wrote the manuscript.

## Acknowledgements

This work was supported by the National Institute of General Medical Sciences and National Institutes of Health through grant R35GM137950 to J.C.R.-M and institutional support from the University of Minnesota Medical School.

